# Cognitive rehabilitation can improve brain injury-induced deficits in behavioral flexibility and impulsivity linked to impaired reward-feedback activity

**DOI:** 10.1101/2023.07.02.547397

**Authors:** Miranda F. Koloski, Christopher M. O’Hearn, Michelle Frankot, Lauren P. Giesler, Dhakshin S. Ramanathan, Cole Vonder Haar

## Abstract

Traumatic brain injury (TBI) affects a large population, resulting in severe cognitive impairments. Although cognitive rehabilitation is an accepted treatment for some deficits, studies in patients are limited in ability to probe physiological and behavioral mechanisms. Therefore, animal models are needed to optimize strategies. Frontal TBI in a rat model results in robust and replicable cognitive deficits, making this an ideal candidate for investigating cognitive rehabilitation. In this study, we report three distinct frontal TBI experiments assessing behavior well into the chronic post-injury period using male Long-Evans rats. First, we evaluated the impact of frontal injury on local field potentials recorded simultaneously from 12 brain regions during a probabilistic reversal learning task (PbR). Next, rats were tested on reversal learning (PbR) or impulsivity (differential reinforcement of low-rate behavior: DRL) and half received salient cues associated with reinforcement contingencies as a form of “cognitive rehabilitation”. After rehabilitation on the PbR task, brains were stained for markers of activity. On the DRL, cues were devalued to determine if beneficial effects persisted on impulsive behavior. TBI resulted in outcome salience deficits evident in task performance and reward-feedback signals occurring at beta frequencies in orbitofrontal cortex (OFC) and associated frontostriatal regions. Cognitive rehabilitation improved flexibility and increased OFC activity. Rehabilitation also reduced impulsivity, even after cues were degraded, which was partially mediated by improvements in timing behavior. The current study established a robust platform for investigating cognitive rehabilitation in animals and identified a strong role for dysfunctional OFC signaling after frontal TBI.

## Introduction

Traumatic brain injuries (TBIs) affect over 2.8 million people each year in the United States and are associated with significant risk for a variety of psychiatric diseases (Vaishnavi et al., 2009). Cognitive deficits, including impaired behavioral flexibility and increased impulsivity also occur at high rates (Sherer et al., 2003; Moreno-López et al., 2016). These deficits are chronic and can substantially affect work productivity, family life, and increase risk for substance abuse and dependence (Miller et al., 2013). Research in cognitive rehabilitation has promise to treat these problems (Novakovic-Agopian et al., 2011; Novakovic-Agopian et al., 2021), but findings are inconsistent due in part to many studies with weaker designs (Fetta et al., 2017). Such issues may have contributed to recent false advertising claims leveraged against commercial “brain training” programs by the US government (Federal Trade Commission, 2016). A stronger understanding of the mechanisms driving impaired behavioral flexibility and impulsivity may yield new insights to develop effective cognitive therapies for TBI.

Behavioral flexibility can be defined as adaptation of behavior in response to changing contingencies (Butts et al., 2013). Flexibility can be measured experimentally by changing reinforcing contingencies after a response strategy is acquired. The Probabilistic Reversal Learning (PbR) task is ideal for repeat testing with its moderately ambiguous reinforcement contingencies (Dalton et al., 2014). Impulsivity is a heterogenous construct which can be defined as actions that may result in short-term gain at the cost of long-term benefits (Ozga et al., 2018). Impulsivity is measured experimentally by various tasks which concurrently pit a reinforced response against the need to inhibit that response under certain conditions (e.g., cues, time). One such task is the differential reinforcement of low-rate behavior (DRL) schedule, which requires a minimum amount of time between responses for reinforcement (Wilson and Keller, 1953).

Both behavioral flexibility and impulsivity are reliant on circuits involving the frontal cortex (medial prefrontal, orbitofrontal cortices) and connections to the dorsal and ventral striatum.

Dopaminergic modulation of these circuits is a critical regulator of such behaviors (Menon et al., 2001; Butts et al., 2013) and striatal tonic and evoked dopamine activity is chronically reduced in a severity-dependent fashion after injury (Chen et al., 2015). These changes in dopamine signaling in frontostriatal networks may lead to reduced outcome salience (i.e., less salient reinforcer-action contingencies) and impair behavioral flexibility or exacerbate impulsivity. Indeed, rats with frontal TBI preferentially track the location associated with reinforcer delivery (i.e., goal-tracking) over reinforcement-predictive cues (i.e., sign-tracking) (Modrak et al., 2020). Thus, a strategy for behavioral rehabilitation may involve augmenting the saliency of reinforcement contingencies and strengthening response-outcome relationships.

The majority of rehabilitative efforts aimed at TBI focus on motor recovery. However, there is strong interest in the emerging area of non-motor, or cognitive, rehabilitation to treat symptoms such as those described above. Effective animal models allow us to study cognitive rehabilitation with greater rigor and gain understanding of the mechanisms by which therapies act. A recent study demonstrated efficacy of spatial-based cognitive rehabilitation for TBI (Law et al., 2023), but was focused on hippocampal-related functions and may not translate for frontally-dependent behaviors such as flexibility and impulsivity.

Thus, the goal of the current study was to establish an animal model for studying cognitive rehabilitation after frontal TBI in rats. We recorded local field potentials (LFP) from 12 frontostriatal regions during the PbR task to better understand the physiology driving post-TBI functional deficits and whether they were location-or frequency-specific. Next, we evaluated a salient environmental cue predicting reinforcement as a form of “cognitive rehabilitation” on the PbR task. We then extended this rehabilitative paradigm to the DRL, using the high-resolution behavioral data to parse impairments in disinhibition from time estimation. To evaluate whether treatment effects persisted, we decoupled cues from reinforcing contingencies and tested for an additional two weeks. Finally, we probed the effects of cues in post-mortem brain using early activation and dopamine markers.

## Materials and Methods

### Experimental Design

In Experiment 1, to study how widespread neural activity is integrated during decision-making, we recorded LFPs while rats performed a self-paced probabilistic learning task (PbR). We used linear mixed models to compare brain activity during reward-feedback collected simultaneously from 12 electrode sites in either TBI or Sham condition animals from weeks 3-11 post-injury (Fig 1A). In Experiments 2 and 3, to determine whether introducing salient cues predicting reinforcing outcomes could rescue chronic TBI-induced impairment, we conducted two behavioral studies, and one histological study. The core design for these studies was a 2 × 2 factorial with animals in either TBI or Sham conditions, and Cue or Control conditions. In Experiment 2, we assessed initial behavioral flexibility using the Attentional Set-Shifting task (AST) and then determined whether impairments could be rescued by cues associated with the highest probability choice on the PbR task (Fig 1B). We then measured neural activity in target regions using histological markers of early activation. In Experiment 3, we used a differential reinforcement of low-rate behavior (DRL)-20 s schedule to measure impulsive behaviors associated with disinhibitory responding and poor discrimination of time intervals. After behavior was stable, cue importance was degraded (always on) to probe whether the intervention had lasting effects on impulsivity (Fig 1C).

**Figure 1.**
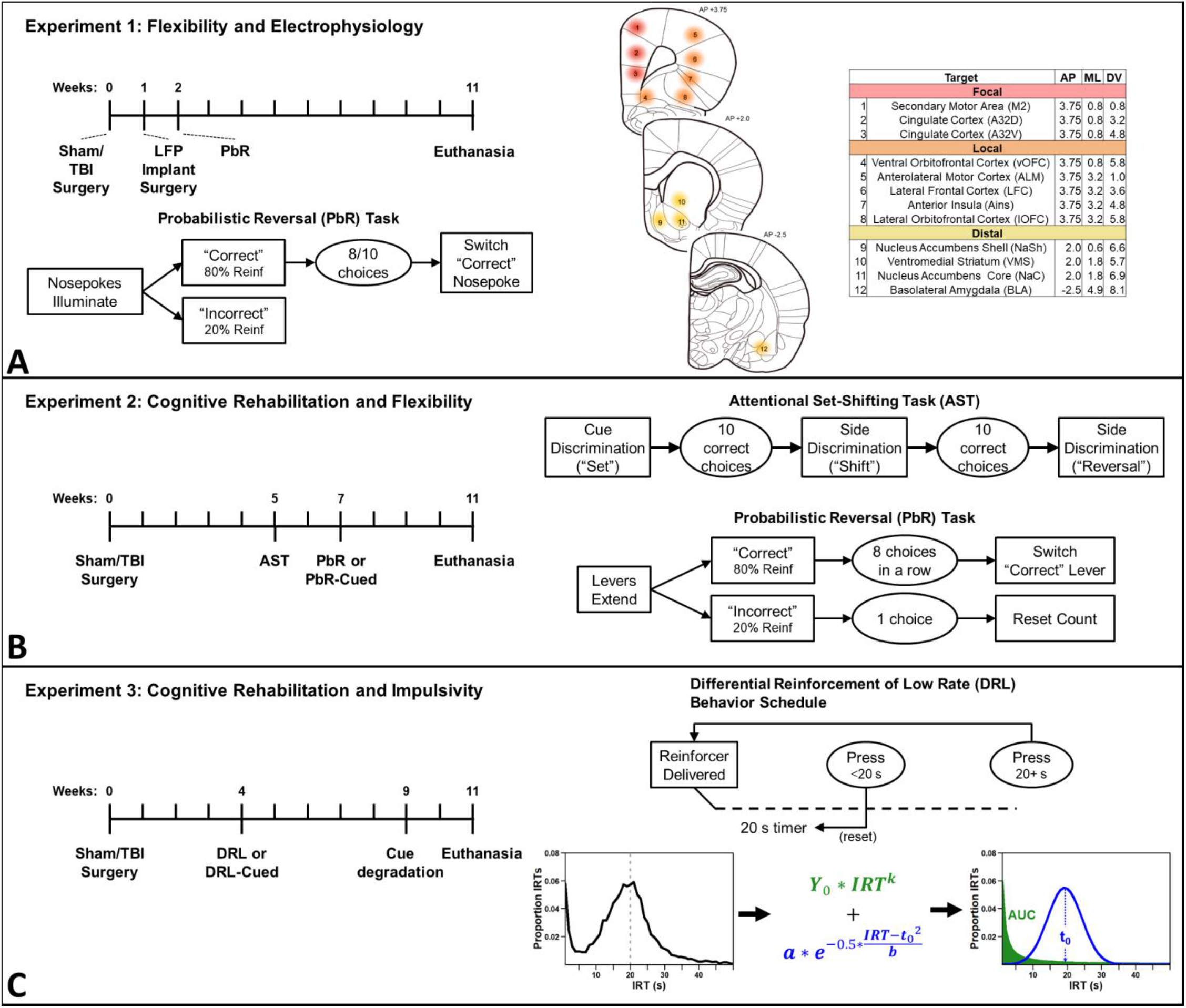
Study design. A) Experiment 1, timeline, task design, and electrode placement. Rats received a TBI, then one week later, electrode implants. Rats were tested on a probabilistic reversal learning (PbR) task starting at two weeks post TBI. PbR was self-paced and used water as a reinforcer. B) Rats received a TBI, then 5 weeks later, tested on the attentional set-shifting task. They were then randomized into Cognitive Rehabilitation or control groups and tested on a PbR task starting at 7 weeks post-surgery. The Rehabilitation condition used a cue light to signal the correct lever. Both used sucrose pellets as reinforcers. C) Rats received a TBI, then 4 weeks later were randomized into Cognitive Rehabilitation or control groups and tested on the differential reinforcement of low rate (DRL) behavior schedule. The Rehabilitation condition used a cue light to signal when 20 s had elapsed. After five weeks, the cue light was turned on for all rats to test for persistence of rehabilitative effects. DRL used sucrose pellets as reinforcers. DRL interresponse time (IRT) data were analyzed by fitting a mixture function consisting of an exponential decay (green) and Gaussian (blue) function to estimate disinhibition (area under curve, AUC) and timing-related impulsivity (blue peak; t_0_ parameter).

### Animals

Male Long-Evans rats (Charles River, Wilmington MA, N = 123) were received at approximately one month old weighing 150g (UCSD, Experiment 1) or approximately 2.5 months of age (WVU, Experiments 2-3). In Experiment 1, rats were pair-housed in standard rat cages (Allentown, NJ, USA) prior to surgery (individually housed following surgery), maintained on a standard light cycle (lights on at 6am/ off at 6pm), and had free access to food. Water was restricted (2 h of free access/ day) during behavioral training to maintain motivation for water reinforcement. Rats were weighed weekly and had free access to water on non-training days. In Experiments 2-3, rats were maintained on a 12:12 h reverse light cycle, with behavioral sessions during the dark cycle and were maintained at approximately 85% *ad libitum* weight and with free water access. Rats were pair-housed in in large pentagonal cages (Optirat, Animal Care Systems) with a divider in the middle. All procedures were conducted in accordance with the National Institutes of Health Guide for the Care and Use of Laboratory Animals and were approved by the West Virginia University Institutional Animal Care and Use Committee or San Diego VA Medical Center Institutional Animal Care and Use Committee (Protocol Number A17-014; A21-012). 7 rats died during surgery, 5 rats were excluded due to lack of damage from the TBI, and 3 were excluded due to lack of electrophysiology signal, thus the final number for the studies was 108. The electrophysiology from sham controls used in Experiment 1 is also expanded upon on in more depth in another publication (Koloski et al., 2023).

### TBI Surgery: Controlled Cortical Impact

Surgical procedures were performed according to previous studies (Martens et al., 2019; Shaver et al., 2019). In brief, each rat was anesthetized with isoflurane (2-4% in 0.5 L/min oxygen) and placed in a stereotaxic frame with a heating pad. The area of incision was cleaned with 70% ethanol and iodine solution and injections of lidocaine were given under the skin for local anesthetic. Rats in Experiment 1 received atropine (0.05 mg/kg, s.c.) to diminish respiratory secretions during surgery and 5-10 mL 0.9% sterile saline. Rats in Experiment 2 received ketoprofen (5 mg/kg, s.c.) prior to surgery and 24 h later and 5 mL 0.9% sterile saline. A 2.0 cm incision was made along the midline to the skull. TBI rats received a 6.0 mm diameter circular craniectomy centered over the prefrontal cortex (AP +3.0, ML +0.0 from bregma) using a micro-drill. A stainless-steel circular impactor tip (5.0 mm diameter) was positioned over the craniectomy and an electromagnetic controlled cortical impactor (Leica Biosystems, Buffalo Grove, IL) was used to induce a focal injury (2.5 mm depth; 3 m/s velocity; 500 ms dwell). The incision was sutured and the rat was placed in a heated recovery chamber until it regained consciousness. Sham rats underwent a similar surgical procedure, except they did not receive a craniectomy or injury. Rats in Experiment 1 received a single dose of buprenorphine SR (1 mg/kg, s.c.) for pain management and were given sulfamethoxazole and trimethoprim in their drinking water (60mg/kg for 5 days) to prevent infection.

### Electrophysiology Implant Surgery

One week following the TBI/Sham procedures, rats in Experiment 1 underwent a second surgery to implant the 32-channel local field potential (LFP) microwires – previously described in detail (Francoeur et al., 2021). Preparatory surgical procedures were the same as described above. After clearing the skull, 8 holes (0.9 mm diameter) were drilled for microwires at predetermined stereotaxic locations. Each hole was for a cannula containing 4 microwires cut to target different DV depths (32 total microwires). An additional hole was drilled above the cerebellum for the ground wire soldered to an anchor screw. Several more holes were drilled for anchor screws (5-8) at the skull periphery. Cannula were lowered to desired depth, pinned to an Omnetics Electrode Interface Board (Neuralynx, MT, USA), secured with Metabond (Parkell, NY, USA), and encased in dental cement (Stoelting, IL, USA). At the conclusion of surgery, the skin was sutured, and rats were given a single dose (1 mg/kg, s.c.) of buprenorphine SR for pain management. Rats recovered from surgery in a heated chamber and received sulfamethoxazole and trimethoprim in their drinking water (60 mg/kg for 5 days) to prevent infections.

### Behavioral Apparatus

For Experiment 1, habituation, pre-training and the PbR task paired with electrophysiology were all performed in custom acrylic operant chambers equipped with five nose ports (each with an LED, IR sensor, and tubing for water delivery), two auditory tone generators, a houselight, and five peristaltic stepper motors to deliver water reward. The chamber was 16 × 12 × 16 inches (L × W × H) with a ceiling opening for electrophysiology tethers. Each operant box was enclosed in a sound-attenuating chamber. Simulink (Mathworks) installed directly onto a Raspberry Pi system controlled the behavioral programs. The design, operation and software control of this chamber has been described previously (Buscher et al., 2020). Electrophysiological data was recorded using a 32-channel RHD head stage coupled to a RHD USB interface board (Intan Technologies, CA, USA) with an SPI interface cable. 12 frontostriatal target electrodes were analyzed in the current study (Fig 1A). Recordings in Open Ephys software were taken at 1KHz with a band-pass filter set at 0.3-999Hz during acquisition. Physiology data was integrated with behavioral streams using a lab-streaming-layer protocol (Ojeda et al., 2014).

Experiments 2 and 3 took place in a bank of 16 standard 5-choice operant chambers (Med Associates, St. Albans, Vermont) enclosed in a sound-attenuating box. Sucrose pellets (Bio-Serv F0021) were used as reinforcers. White noise generators were used to further mitigate any effects of extraneous sounds. The right panel of each chamber was equipped with a food hopper and two retractable levers on either side. White stimulus-lights were above the levers and a houselight was at the top of the chamber. The 5-choice array was located on the opposite side and was not used in the current study. Custom programs were written (DRL) or adapted from Dr. Stan Floresco (AST, PbR) using MedPC software.

### Experiment 1: Electrophysiological Correlates of Behavioral Flexibility

This experiment evaluated reward-feedback related neural activity following TBI associated with behavioral flexibility performance on the PbR task. Prior to surgery, animals were habituated (three sessions) and trained to perform sequential nose port responses for water reinforcers but were naïve to the final PbR task. Animals began habituation 7 months after arrival. On average, training took two weeks to complete (5-15 sessions), at which time surgeries were conducted. Once rats recovered from the second surgery, they began training on the self-paced PbR task (two weeks post-injury) (Fig 1A). The behavioral training and simultaneous electrophysiology recording occurred in 60-minute sessions two-four times/week in Sham (n= 10) and TBI (n = 12) rats.

On the PbR task, different probabilities of reinforcement were randomly assigned to each choice port at the start of a session (“correct” nose port = 80% reinforcement, “incorrect” nose port = 20% reinforcement). Each trial began with houselights off and the middle nose port LED on. Responses to the middle nose port initiated trials. LEDs in the choice ports to the left and right of the middle nose port to illuminated to signal an available choice. A response in the “correct” port led to 2 s (20 μL) of water 500 ms after the response on 80% of trials. Selecting the “incorrect” port led to 2 s (20 μL) of water on only 20% of trials. Trials with no reinforcer resulted in tone presentation (500 ms) to signal lack of water and illumination of the houselight. There was a 5-s intertrial interval after the outcome. The probabilities associated with each choice port were reversed when eight out of the last ten responses in a moving window were “correct” choices (regardless of reinforcement outcome). For rats with implants still intact (n = 17), the experiment ended at 11 weeks post-injury (minimum included was 8 weeks post-injury). Analyses were based on 275 behavioral sessions (average 11 sessions/ rat). See Figure 1A.

After the experiment, rats were anesthetized with a lethal dose of isoflurane and euthanized by transcardiac perfusion with 0.9% phosphate buffered saline, followed by 4% phosphate buffered formaldehyde. Brains were post-fixed in 4% phosphate buffered formaldehyde for 24 h before being transferred to a 30% sucrose solution. Tissue was blocked in the flat skull position in 3mm coronal sections which were paraffin embedded and sectioned 20 μm thick on a microtome. Cut tissue was floated in a 40°C hot water bath and mounted on slides. Slices were deparaffinized and stained with thionin to visualize cell body loss from injury and map electrode tracks. Sections were processed with a slide scanner at 40x magnification (Zeiss, Oberkochenn, Germany; Leica Biosystems, IL, USA).

LFP data was pre-processed offline using custom MATLAB scripts and functions from EEGLAB (Fakhraei et al., 2021b; Fakhraei et al., 2021a; Francoeur et al., 2021). Data was aligned to the choice response to examine neural activity during the reward-feedback period (500 ms to 2500 ms after response). For artifact removal, activity was averaged across time and electrodes to compute a single value for each trial. Trials with a value greater than 4X standard deviation were treated as artifact and removed. The remaining trials were median referenced and a trial-by-trial time-frequency decomposition was calculated using a complex wavelet function in EEGLAB (*newtimef* function, using Morlet wavelets). For each frequency and electrode, we subtracted the mean activity within a baseline window (1000-750 ms prior to the start of the trial) to calculate the evoked power change from baseline. For each session, activity was averaged across trials for each choice and reward outcome (“correct” reward, “correct” no reward, “incorrect” reward, “incorrect” no reward).

### Experiment 2: Cognitive Rehabilitation and Behavioral Flexibility

This experiment evaluated common paradigms for assessing behavioral flexibility and implemented a salient cognitive rehabilitation intervention to improve function after TBI, then evaluated neural activity in post-mortem tissue using immediate early gene markers.

### Experiment 2a: Attentional Set-Shifting

To establish a baseline for flexibility deficits prior to long-term testing, the AST was carried out as previously described (Butts et al., 2013), at 5 weeks post-injury in Sham (n = 22) and TBI rats (n = 19). Rats were habituated (two sessions, 10 sucrose pellets freely available) to the chamber and then trained to lever press using an autoshaping procedure where a pellet was delivered every 35 s on average, but 10 s prior to pellet delivery, both levers extended, and any presses were immediately reinforced. Three sessions were conducted until 40 presses occurred, or pressing was hand shaped by the experimenter. Rats then underwent retractable lever press training and a side preference assessment. AST testing began the next day. Each phase of AST consisted of 200 discreet trials per day, or until 10 consecutive correct responses occurred. The initiation of a trial was marked by the illumination either the left or right stimulus-light which varied pseudorandomly. After 3 s, both the left and right levers were extended, and the rat was allotted 10 s to make a response. Phase 1 (“set”), the cue discrimination task, reinforced responses to the location of the stimulus light. Phase 2 (“shift”), the response discrimination task, reinforced responses to one side (left/right), regardless of light position. Phase 3 (“reversal”), the response reversal task, reinforced responses to the side opposite the previous phase (e.g. left to right). See Figure 1B.

### Experiment 2b: Probabilistic Reversal Learning and Cognitive Rehabilitation

One week after completing the AST, rats began PbR testing and continued for 5 weeks according to previous protocols (Dalton et al., 2014). Each PbR session consisted of 200 discrete trials separated by a 15-s intertrial interval; trials began on presentation of both levers and lasted 10 s. During the first trial, differential probabilities of reinforcement were assigned to each lever (“correct” lever = 80%, “incorrect” lever = 20%) as described in Experiment 1. After eight consecutive “correct” choices (regardless of reinforcement), the probabilities associated with each lever reversed. See Figure 1B. To model cognitive rehabilitation, approximately half the rats had a cue light illuminated above the “correct” lever 3 s prior to extension and throughout the trial, resulting in four groups: Sham (n=11), TBI (n=10), Sham-Cue (n=10), and TBI-Cue (n=9). Rats were matched for AST performance and pseudorandomly assigned to cue or control condition.

### Experiment 2c: Histology

During week 11 post-injury, and at 90 minutes after behavioral testing, rats were anesthetized with a lethal dose of pentobarbital and transcardially perfused with ice-cold 0.9% phosphate buffered saline, followed by 3.7% phosphate buffered formaldehyde. Brains were post-fixed in 3.7% phosphate buffered formaldehyde for 24 h before being transferred to a 30% sucrose solution. Once fully saturated, the brains were embedded into gel blocks (15% gelatin; 4-5 brains per block), post-fixed in 3.7% phosphate buffered formaldehyde for 24 h, then transferred to a 15% sucrose solution for two days, and then a 30% sucrose solution until fully saturated. Blocks were then frozen and sliced on a microtome at 30 μm along the coronal plane.

Individual slices were then stained with antibodies against c-Fos, to visualize cells active during behavioral testing, and tyrosine hydroxylase (TH), to visualize cells producing dopamine. Slices were blocked in 5% normal goat serum overnight at 4°C, then co-incubated in anti-c-Fos (ab208942; 1:500) + anti-TH (ab112; 1:1000) for 72 h at 4°C. Secondary antibodies were then applied in separate overnight incubations: anti-Mouse IgG (ab150116; 1:2000) and anti-Rabbit IgG (ab150077; 1:2000).

Slides were then imaged on an Olympus BX-43 with a DP-80 12MP camera in several regions of interest: prelimbic (PL), orbitofrontal cortex (OFC), and nucleus accumbens core (NAc). One image was taken at 10x magnification for each region, then split into quadrants for quantification. ImageJ (NIH, Bethesda, MD) was used to manually count c-Fos^+^ cells after thresholding the image. The c-Fos (red) and TH (green) images were then merged using ImageJ’s Colocalization macro. Because dopamine cells were sparse outside of the NAc and because dopaminergic projections to frontal regions were also of interest, any cell with multiple colocalized pixels clustered together was counted as putative dopaminergic synapses onto active cells marked with c-Fos.

### Experiment 3: Cognitive Rehabilitation and Impulsivity

The following experiments evaluated whether cognitive rehabilitation could rescue impulsive deficits and if they persisted after rehabilitation was removed.

### Experiment 3a: Differential Reinforcement of Low-Rate Behavior

Two weeks post-injury, rats were habituated and underwent autoshaping as described in Experiment 2, but with only the left lever extended. Three sessions were conducted until 40 presses occurred, or pressing was hand shaped by the experimenter. Rats then performed 3 sessions of fixed ratio-1 responding. At 4 weeks post-injury, rats began testing on 60-minute sessions of DRL-20 for 5 weeks. On the DRL, presses were only reinforced if they were spaced by greater than 20 s. Any press made prior to that reset the timer (Wilson and Keller, 1953). See Figure 1C. To evaluate cognitive rehabilitation, approximately half the rats had a cue light illuminated after the 20 s timer had elapsed, resulting in four groups: Sham (n=13), TBI (n=10), Sham-Cue (n=11), and TBI-Cue (n=11).

### Experiment 3b: Persistence of Cognitive Rehabilitation Effects

After 5 weeks of DRL performance, behavior was stable. To determine whether the cue training persistently improved impulse control, the salience of the cue was degraded by turning on the cue light at all times for all rats. Rats were then re-tested for two additional weeks.

### Statistical Analysis

The primary behavioral outcome measure for the PbR task in Experiment 1 was the number of reversals completed. Secondary measures included total number of trials, initial discrimination trials (trials before first reversal), and likelihood of switching given a win or loss. The primary outcome measures for Experiment 2 on the AST were the trials to criterion for each phase, on the PbR number of reversals completed, and on the histology the number of c-Fos^+^ cells and the percent of colocalized cells. Secondary measures included omitted trials (AST, PbR) and likelihood of switching given a win or loss (PbR) and errors to criterion (AST). The primary outcome measures for Experiment 3 on the DRL were the percent of reinforced responses and the distribution of interresponse times (IRTs). Total responses served as a secondary measure.

In Experiment 1, LFP data was first analyzed across frequency bands of interest (delta (1-4Hz), theta (4-8 Hz), alpha (8-12 Hz), beta (15-30 Hz), low-gamma (40-70Hz), and high-gamma (70-150Hz) power) at the lateral orbitofrontal cortex (lOFC) electrode. Electrophysiology data were analyzed using linear mixed-effects models (LMER; normalized power) using frequency band as the slope parameter of the random effect for each subject (Frequency|Subject) and Time, and Injury (Sham/TBI), Frequency (delta, theta, alpha, beta, low-gamma and high-gamma), Trial (Reward/ No Reward) and their interactions as the fixed effects. Frequency band was assigned to the slope parameter of the random effect for each subject to account for the interdependence of power at each frequency band (Frankot et al., 2023). The estimated marginal means were plotted separately for Reward trials, No Reward trials, and their contrast [Reward-No Reward]. Next, the normalized power at 12 electrode locations focal, local, and distal to the injury were analyzed at frequency bands with relevant between-group (Sham/TBI) differences. Another linear mixed-effects model was used to examine normalized power (LMER; normalized power) using individual Subject intercepts and Time as the random effect, and Injury (Sham/TBI) and Electrode (12) and their interactions as the fixed effects. The estimated marginal means were plotted separately for Reward trials, No Reward trials, and their contrast [Reward-No Reward]. The critical *p*-value was set at 0.05. Post-hoc tests were Bonferroni corrected. All data were analyzed using SPSS and visualized using Graph Pad Prism software.

Transformations were applied as necessary to normalize data. A Box-Cox test was used to determine transformations. Transformations included log for variables bounded at the lower end (AST trials/errors to criterion, PbR reversals, histology c-Fos^+^ counts), square root for skewed variables (DRL percent reinforced, histology percent colocalized), inverse square root (DRL total responses, PbR omissions), inverse (AST omissions), square (PbR shift probabilities). The critical *p*-value was set to 0.05. All data were analyzed using R statistical software (http://www.r-project.org/; version 4.0.2) with the *lme4, lmerTest* and *stats* libraries.

Most repeated-measures behavioral data were analyzed using linear mixed-effects regression (LMER; DRL percent reinforced, PbR reversals, omissions, and stay probability) using individual subject intercepts as the random effect, and Injury (Sham/TBI), Rehab (Non-Cued/Cued), Week and their interactions as the fixed effects. The AST data were analyzed with repeated measures ANOVA (Injury × Phase) because there was only one observation per phase per subject, and post-hoc tests were conducted with a Student’s t-test since there were only two groups. A planned comparison was made with a t-test for the last session of performance in Experiment 3b between the TBI and TBI-cue group to determine if rehabilitation had a lasting beneficial effect. Histological data were analyzed as a mixed-effects ANOVA with individual subject as the random effect and Injury, Rehab, and their interactions as fixed effects.

IRT distributions in the DRL were analyzed by fitting a compound curve (using *nls* in R) consisting of an exponential decay and gaussian distribution (Fig 1C). From this function, disinhibitory responding was calculated by taking the area under the curve (AUC) up to the 5 s point and the ability to time the delay were estimated from the *t*_*0*_ value of the equation. These values were then used as outcome measures in LMERs as described above. Finally, a planned comparison was made with a t-test for the last session of performance in Experiment 3b between the TBI and TBI-cue group to determine if rehabilitation had a lasting beneficial effect. Prior studies have commonly divided IRTs into early and late (Cheng et al., 2006; Fowler et al., 2009), but this fails to model the inter-dependency between the two distributions (i.e., since they are proportions, change in one parameter necessitates changes in others) which we have recently demonstrated drastically affects false positive rates (Frankot et al., 2023).

## Results

### Brain Injury Impaired Probabilistic Reversal Learning

All outcome variables on the PbR task were analyzed using LMERs [Fixed: Injury (Sham, TBI) x Time, Random: Subject]. For the primary outcome measure, the number of reversals, the interaction of Injury × Time was significant (*p* = 0.001; Fig 2A; see Supplemental Table 2–1 and 2–2). TBI rats performed fewer reversals than Sham rats and had a slower rate of learning (*p* < 0.001). Since the task was self-paced, the lower rate of reversals may have been mediated by the total number of trials completed, however injury did not significantly change trial count between groups (Injury *p* = 0.055, Fig 2A). On average, TBI animals completed 86.69 (±15.62) trials and Sham animals completed 134.76 (±15.50) trials with both groups increasing the number of trials across time (Time *p* < 0.001, Fig 2A).

**Figure 2.**
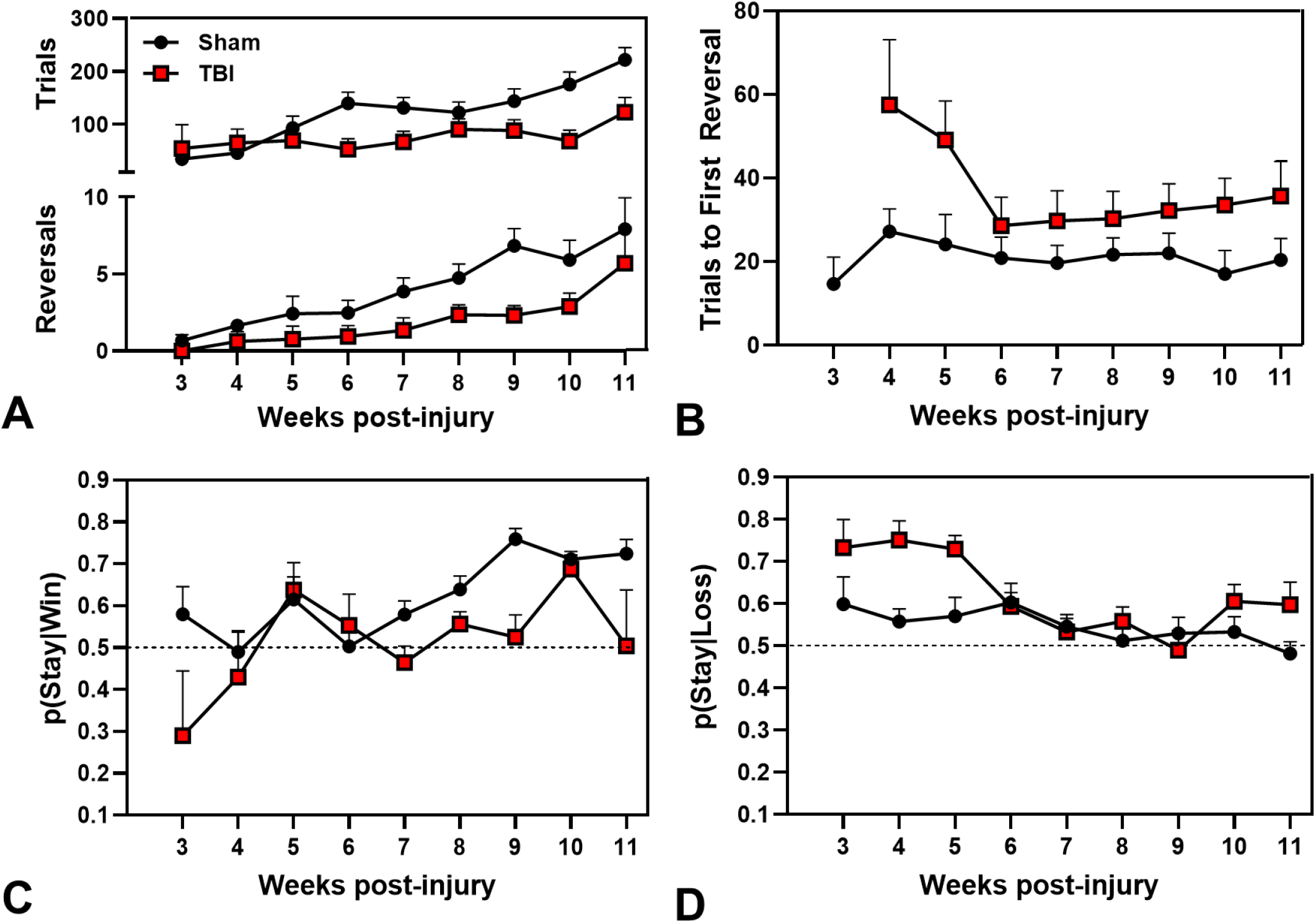
Experiment 1: Performance on the self-paced PbR task. A) TBI (red square) significantly reduced the number of reversals compared to Sham (black circle; *p* < 0.001), with deficits persisting 11 weeks after injury, but there were no differences in trials between groups (*p* = 0.055). B) TBI rats required more trials to complete the first discrimination block *(p* < 0.001). There is no TBI week 3 data point because rats did not meet criterion for a reversal. C) After a winning trial, rats were more likely to stay with the same option over time (*p* < 0.001), but there was no significant difference in Win-Stay behavior between groups. D) After a losing trial, Sham rats were less likely to choose the same option relative to TBI (*p* = 0.016), and both groups showed less Lose-Stay behavior over time (*p* = 0.002). Symbols represent group estimated marginal means (+SEM).

To more closely examine reward learning deficits related to behavioral flexibility, we assessed the number of trials in the initial discrimination block (Fig 2B) and the probability of choosing the same option given a win ([p(Stay|Win)]; Fig 2C) or a loss ([p(Stay|Loss)]; Fig 2D). TBI rats took more trials to complete the first reversal (initial discrimination block) compared to Sham rats (*p* < 0.001; Fig 2B). There was no difference between groups in Win-Stay behavior (Fig 2C). However, the probability of choosing the same option after a loss (no reward) was greater in TBI than Sham groups (*p* = 0.033; Fig 2D). There was also a main effect of Time (*p =* 0.002; Fig 2D) such that the probability of making the same choice after a loss decreased across time.

### Beta Power was Modulated by Reward Outcome and Impaired after TBI

First, to identify frequency bands of interest during reward-feedback, we ran a linear mixed model to examine normalized power across frequencies on the lOFC electrode [Fixed: Injury (Sham, TBI) × Trial (Reward, No Reward), Frequency (Delta, Theta, Alpha, Beta, LGamma, HGamma). Random: (Frequency|Subject), Time]. Trial Type and Frequency were repeated measures. We selected the lOFC electrode because it is a cardinal reward-processing region in the frontostriatal network and is located outside the focal injury site. There was a main effect of Trial (Reward, No Reward) (*p* < 0.001; Fig 3A), and main effect of Frequency (*p* = 0.006; Fig 3A; see Supplemental Table 3–1). There were also significant interactions between Injury x Trial (*p* < 0.001; Fig 3A) and Trial x Frequency (*p* < 0.001; Fig 3A). Post-hoc comparisons reveal that lOFC power was greater on Reward (0.73 ± 0.12) compared to No Reward (0.06 ± 0.12) Trials. On Reward Trials, Sham rats had significantly greater beta frequency power (*t*_(64.81)_ = 3.07, *p* = 0.019) and greater delta frequency power (*t*_(64.81)_ = 2.91, *p* = 0.029; Fig 3A) compared to TBI rats. On No Reward Trials, there was no difference in power between Sham and TBI animals at any frequency band (Fig 3A). TBI and Sham rats had the largest difference in power on [Reward-No Reward] Trials at beta frequencies (Mean Difference = 2.76 ± 0.85; Fig 3A). Beta activity at the lOFC electrode clearly differentiates between reward outcomes in Sham rats, but the difference is attenuated after TBI when plotted across the reward-feedback period (Fig 3B).

**Figure 3.**
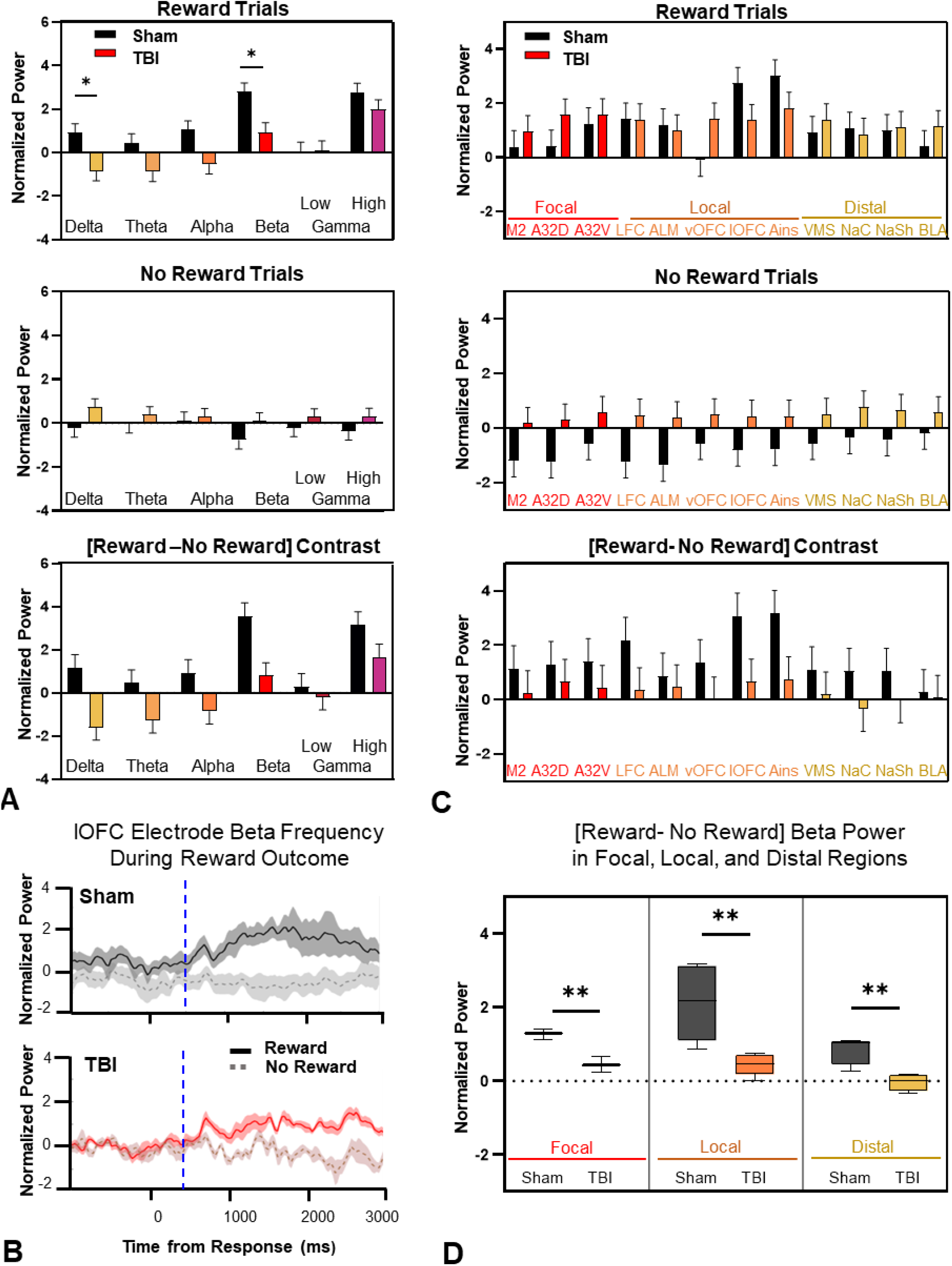
Experiment 1: Local field potential activity during reward outcome of the PbR task. A) Normalized power on the lOFC electrode during reward outcome (500-2500ms after the response) across 6 frequency bands: Delta (1-4Hz); Theta (4-8Hz); Alpha (8-12Hz); Beta (15-30Hz); Low-Gamma (40-70Hz); High-Gamma (70-150Hz). On Reward trials, TBI rats had decreased delta (*p* = 0.029) and beta power (*p* = 0.019) compared to Sham rats. On No Reward trials, there was no significant difference. The difference in power on [Reward-No Reward] Trials was greatest for beta oscillations (*p* = 0.011). Bars represent estimated marginal mean (+SEM) of Sham rats (black) and TBI rats (colorful). B) Beta frequency power on the lOFC electrode during reward outcome is increased on Reward and decreased on No Reward trials for Sham rats. The lOFC beta power difference is attenuated in TBI rats. Mean and SEM (shaded error) traces for Reward (solid) and No Reward (dashed) Trials are time-locked to response (0 ms). The vertical blue line represents the onset of reward delivery. C) Since beta frequency oscillations showed the greatest disruption after TBI, we analyzed 12 electrode sites that were focal (red), local (orange), and distal (yellow) to the injury location. Beta power was greater on Reward trials (*p* < 0.001) and TBI rats showed diminished reward-related beta activity across most electrodes (*p* < 0.001). Post-hoc t-tests confirmed differences in reward-related beta power between TBI and Sham rats at lOFC (*p* = 0.011) and Ains (*p* = 0.037) electrodes. Injury increased beta power on No Reward trials relative to Sham on A32D (*p* = 0.024), LFC (*p* = 0.007), ALM (*p* = 0.013), and lOFC (*p* = 0.030) electrodes. Bars represent estimated marginal mean (+SEM) of Sham rats (black) and TBI rats (colorful). Focal: M2 (secondary motor area); A32D (dorsal cingulate cortex); A32V (ventral cingulate cortex); Local: LFC (lateral frontal cortex); ALM (anterolateral motor cortex); vOFC (ventral orbitofrontal cortex); lOFC (lateral orbitofrontal cortex); Ains (anterior insula); Distal: VMS (ventromedial striatum); NaC (nucleus accumbens core); NaSh (nucleus accumbens shell); BLA (basolateral amygdala). D) Estimated marginal means of beta power averaged across electrodes in focal, local, and distal regions show the difference between [Reward-No Reward] trials are less following TBI (color) compared to Sham rats (grayscale). TBI rats have a smaller difference in beta power between Trials on focal (*p* = 0.004), local (*p* = 0.008), and distal (*p* = 0.008) electrodes compared to Sham. Box and whisker plot show the lower and upper quartiles, mean, minimum and maximum. ^**^ *p* < 0.01; ^*^ *p* < 0.05.

To further examine reward-feedback signals following TBI, we used a linear mixed model to analyze differences between Injury groups across 12 different electrodes in the frontostriatal network. These 12 electrodes were selected to include areas within the focal site of brain injury (M1, A32D, A32V) areas local to the injury site (LFC, ALM, vOFC, lOFC, Ains) and distal sites that receive dense projections from prefrontal cortex related to decision-making (VMS, NaC, NaSh, BLA; Fig 1A). The LMM [Fixed: Injury (Sham, TBI) × Electrode (12 electrodes) × Trial (Reward, No Reward). Random: Subject, Time] with Electrode and Trial analyzed as repeated measures, revealed a main effect of Electrode (*p* < 0.001; Fig 3C), and main effect of Trial (*p <* 0.001; Fig 3C; see Supplemental Table 3–1). There were significant interactions between Injury × Electrode (*p =* 0.013; Fig 3A), Injury × Trial (*p <* 0.001; Fig 3A), and Trial × Electrode (*p <* 0.001, Fig 3A). On Reward Trials, post-hoc tests revealed that TBI rats had significantly lower beta power on lOFC (*t*_(47)_ = 2.39, *p* = 0.011) and Ains electrodes (*t*_(47)_= 2.17, *p* = 0.037). On No Reward Trials, TBI rats showed an increase in beta power relative to Sham. Post-hoc tests revealed a significant difference between groups on the A32D (*t*_(49)_ = -2.33, *p* = 0.024), LFC (*t*_(49)_ = -2.82, *p =* 0.007), ALM (*t*_(49)_ = -2.58, *p = 0*.013), and lOFC electrodes (*t*_(49)_ = -2.23, *p =* 0.030). On average, Sham rats had a larger difference in beta power between [Reward-No Reward] Trials (1.48 ± 0.77) than TBI rats (0.26 ± 0.74) (*p <* 0.001; Fig 3C), which was most robust on the lOFC and Ains electrodes. Although there was a significant effect of Electrode, grouping regions into focal, local, or distal categories revealed the widespread consequence of injury to reduce the difference between reward outcomes (Focal *t*_(4)_= 5.84, *p* = 0.004 ; Local *t*_(8)_ = 3.53, *p* = 0.008; Distal *t*_(6)_ = 3.85, *p* = 0.008 (Fig 3D). The above illustrates a reward-feedback signal occurring at beta frequencies that differentiates between reward outcomes and is impaired following TBI.

### Cognitive Rehabilitation Rescued Deficits in Probabilistic Reversal Learning

Prior to cognitive rehabilitation, rats were tested on the AST. There were no TBI effects, other than a small increase in omitted trials in the reversal phase (*p* = 0.042; see Supplemental Fig 4–1 and Table 4–2). Rats were then pseudorandomly assigned (based on AST performance) to cognitive rehabilitation or normal conditions on the PbR task (see Fig 1B). All outcome variables on the PbR task were analyzed using LMERs [Fixed: Injury (Sham, TBI) × Rehab (Non-Cued, Cued) × Time, Random: Subject; see Supplemental Table 4–3 and 4–4].

For the primary outcome measure, the number of reversals, the 3-way interaction of Injury × Rehab × Time was significant (*p* = 0.042; Fig 4A). Evaluation of this effect revealed that because TBI rats started lower than Shams, they had higher rates of learning (*p* = 0.026). In contrast, cognitive rehabilitation restored reversals in TBI-Cue rats to Sham levels from the start of testing and had no difference in learning rate (*p* = 0.473). For total omissions on the PbR task, there was an Injury × Time effect (*p* < 0.001) such that TBI rats started high but declined over time (Fig 4B).

**Figure 4.**
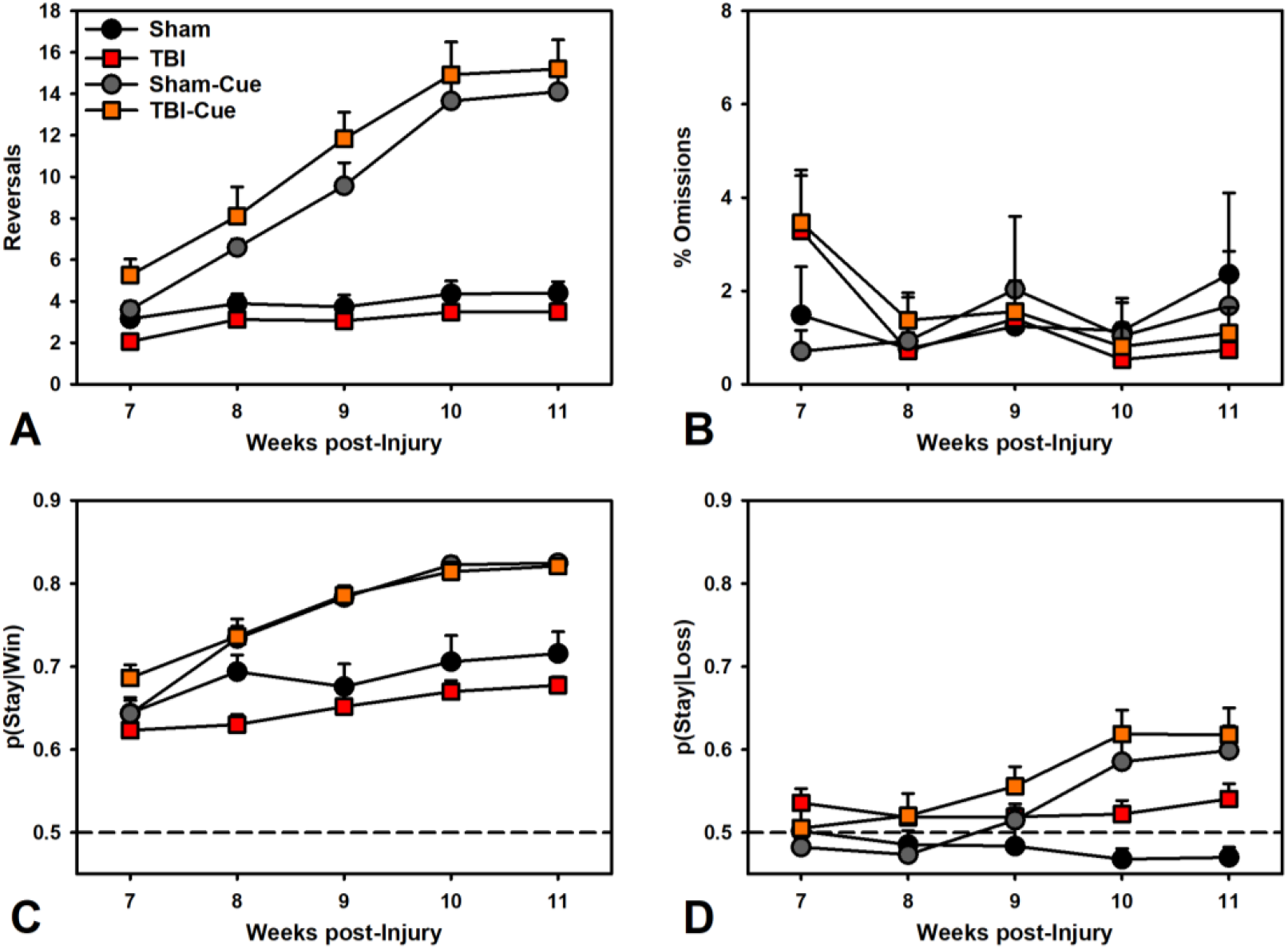
Experiment 2: Performance on the PbR task. A) TBI significantly reduced reversals compared to Sham (*p* = 0.026), while cognitive rehabilitation restored function in TBI-Cue rats compared to Sham-Cue (*p* = 0.473). B) TBI caused an initial small increase in omissions which decreased to sham levels over time (*p* = 0.003). C) After a winning trial, Sham rats were more likely to stay with the same option over time relative to TBI (*p* = 0.008), and cognitive rehabilitation also increased this likelihood (*p* < 0.001). D) After a losing trial, Sham rats were less likely to choose the same option relative to TBI (*p* = 0.029), but cognitive rehabilitation increased likelihood of staying with the same option over time (*p* < 0.001). Symbols represent group means (+SEM).

To better understand the pattern of choice, the probability of choosing the same option given a win [p(Stay|Win)] or a loss [p(Stay|Loss)] were examined (Fig 4C-D). For staying on wins, there was a significant Rehab × Time effect (*p* < 0.001), such that Cued rats had increased likelihood of choosing the same option again relative to Non-Cued rats. There was also a significant Injury × Time effect (*p* = 0.041), such that TBI decreased the likelihood of choosing the same option again relative to Sham. For staying on losses, there was a significant effect of Rehab × Time (*p* < 0.001), such that Cued rats had increased likelihood of choosing the same option again relative to Non-Cued rats. There was also a main effect of Injury (*p* = 0.012) such that TBI rats were more likely to stay after a loss.

### Cognitive Rehabilitation Increased c-Fos Expression in the Orbitofrontal Cortex

To evaluate how cognitive rehabilitation and reversal learning were affecting neural activity, rats were euthanized after completing the task and examined for c-Fos expression. The number of c-Fos^+^ cells were analyzed in mixed-effects Poisson regressions [Fixed: Injury (Sham, TBI) x Rehab (Non-Cued, Cued), Random: Subject; see Supplemental Table 5–1] and c-Fos^+^ cells colocalized with TH^+^ were analyzed in two-way ANOVAs [Injury x Rehab].

For c-Fos^+^ cells, there were no differences in the PL or NAc (*p*’s > 0.107). However, in the OFC, there was a significant Injury x Rehab interaction (*p* = 0.006; Fig 5B), such that TBI severely reduced the number of active cells, but cognitive rehabilitation improved this. For TH-colocalized cells, there was a significant main effect of Injury (*p* = 0.039; Fig 5A), such that TBI increased c-Fos^+^/TH^+^ cells. There were no other effects for colocalized cells (*p*’s > 0.214).

**Figure 5.**
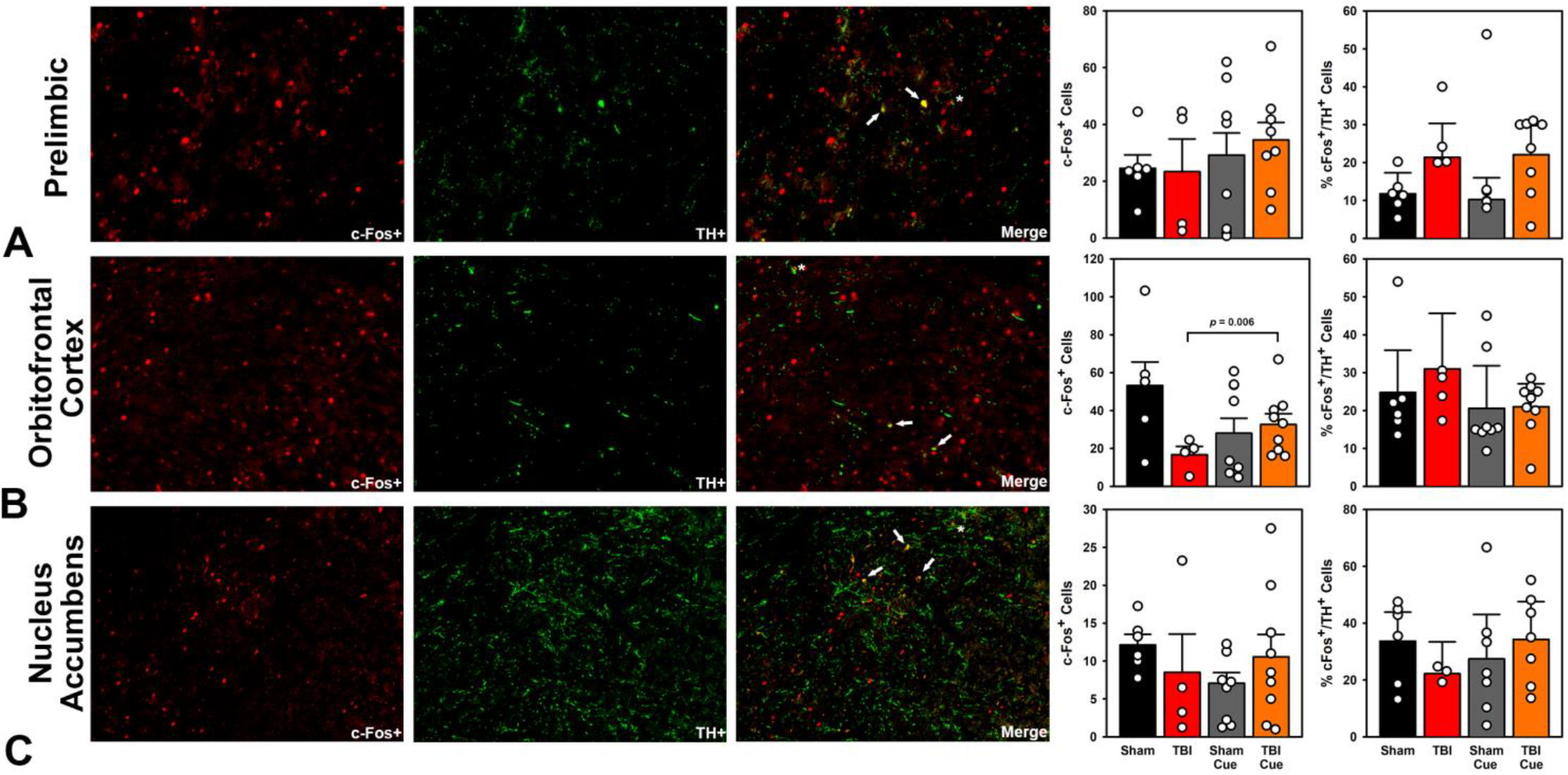
Experiment 2: Active neurons following the PbR task. Panel A shows prelimbic neurons, Panel B shows OFC neurons, and Panel C shows nucleus accumbens core neurons. From left to right in each panel are c-Fos^+^ cells (red), TH^+^ cells (green) and their merge (yellow), followed by quantification of c-Fos+ cells and percent of c-Fos cells colocalized with TH. TBI increased the percent of TH+ active cells in PL (*p* = 0.039). Cognitive rehabilitation increased active OFC cells in TBI rats (*p* = 0.006). Bars represent means (+SEM) and dots represent individual subjects. The ← identifies active dopamine cells, while the ^*^ identifies putative dopaminergic synapses onto active cells.

### Cognitive Rehabilitation Improved DRL Performance after TBI

Percent reinforced responses were analyzed using LMER [Fixed: Injury (Sham, TBI) x Rehab (Non-Cued, Cued) x Time, Random: Subject; see Supplemental Table 6–1 and 6–2]. The 3-way interaction of Injury × Rehab × Time was significant (*p* < 0.001; Fig 6A). A breakdown of this effect revealed that injury impaired TBI rats relative to Sham over time (*p* < 0.001), but cognitive rehabilitation led to no difference between TBI-Cue and Sham-Cue rats (*p* = 0.309). Because these data were percentage-based, a secondary measure of total responses was also analyzed in the same fashion to verify effects were not driven by low response rates in the TBI rats. The 3-way interaction of Injury × Rehab × Time was significant (*p* < 0.001). Comparison of this effect revealed the same findings: that injury impaired TBI rats relative to Sham over time (*p* < 0.001), but there was no difference between TBI-Cue and Sham-Cue rats (*p* = 0.484). Only six instances of less than 100 presses were recorded (lowest: 65), and the average was 260.64 presses per session (data not shown).

**Figure 6.**
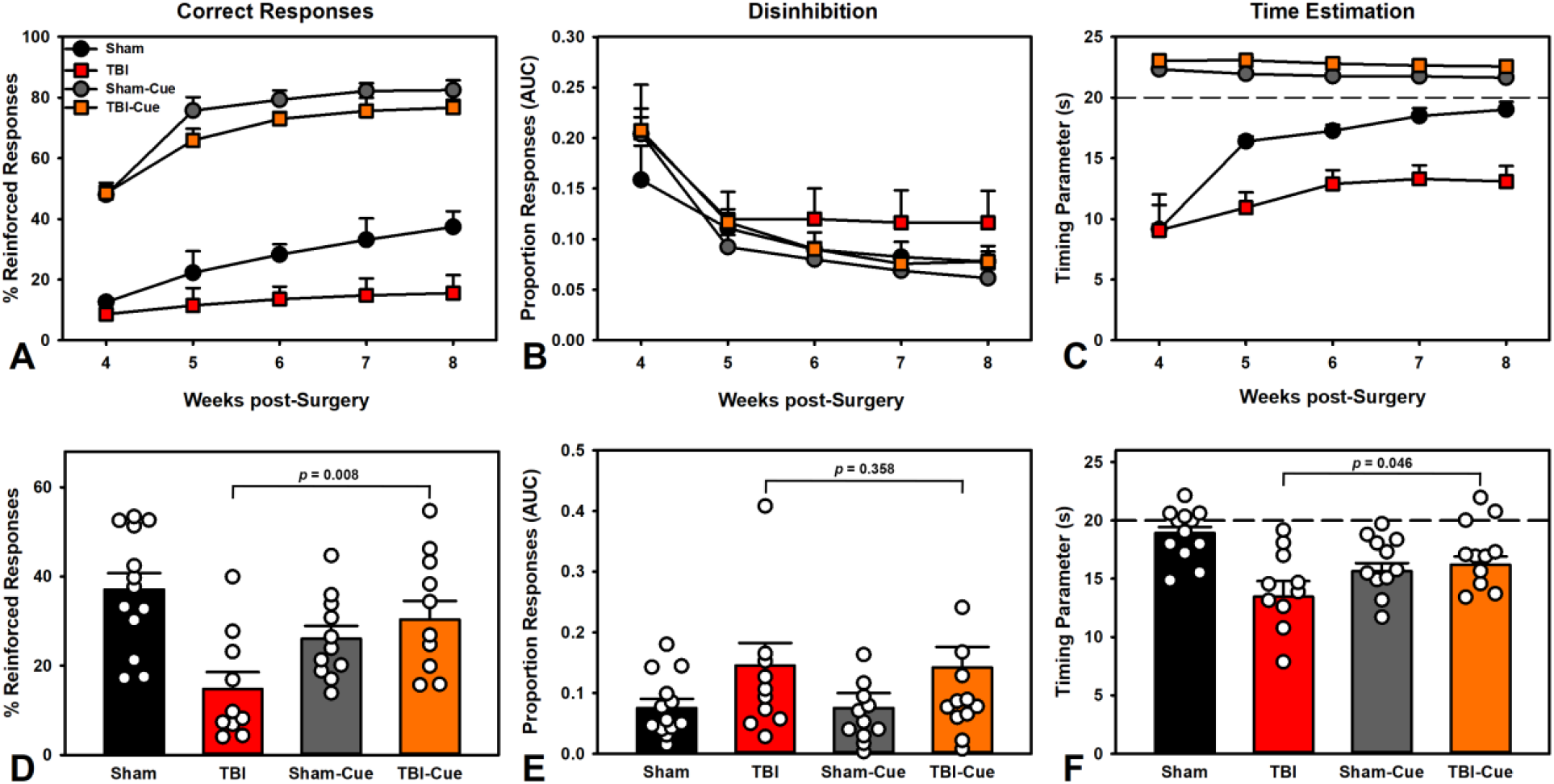
Experiment 3: Performance on the DRL task (A-C) and persistence of cognitive rehabilitation (D-F). A) Sham rats made significantly more correct responses than TBI rats (*p* < 0.001), but cognitive rehabilitation eliminated these deficits in TBI-Cue rats (*p* = 0.253). B) Cognitive rehabilitation reduced overall disinhibited responding (*p* = 0.026), but there were no TBI-specific effects. C) TBI impaired timing estimation compared to Sham (*p* < 0.001), but cognitive rehabilitation eliminated this deficit in TBI-Cue rats (*p* = 0.968). D) Cognitive rehabilitation improved correct responding in TBI-Cue animals even after cues were devalued (*p* = 0.008). E) This was not due to reduced disinhibition (*p* = 0.358). F) However, this improvement was partially accounted for by improvements in timing (*p* = 0.046). Symbols and bars represent group means (+SEM) and dots represent individual subjects.

Individual IRT distributions were fit to the compound exponential + Gaussian curve (see Fig 1C) using one week’s worth of data per subject. The raw curves are shown in Figure 7. Parameter values were then used to estimate proportion of disinhibitory responding using the AUC and ability to time the delay using the t_0_ parameter. These values were analyzed in a LMER as described above. For disinhibitory responding, only the Rehab × Week effect was a significant contributor to the model (*p* = 0.023; Fig 6B) such that the Rehab reduced disinhibitory responses. For timing, the 3-way interaction of Injury × Rehab × Time was significant (*p* = 0.006; Fig 6C); TBI rats had significantly left-shifted values (i.e., shorter IRTs) over time relative to sham (*p* < 0.001), while cognitive rehabilitation prevented this effect in TBI-Cue rats relative to Sham-Cue (*p* = 0.968).

**Figure 7.**
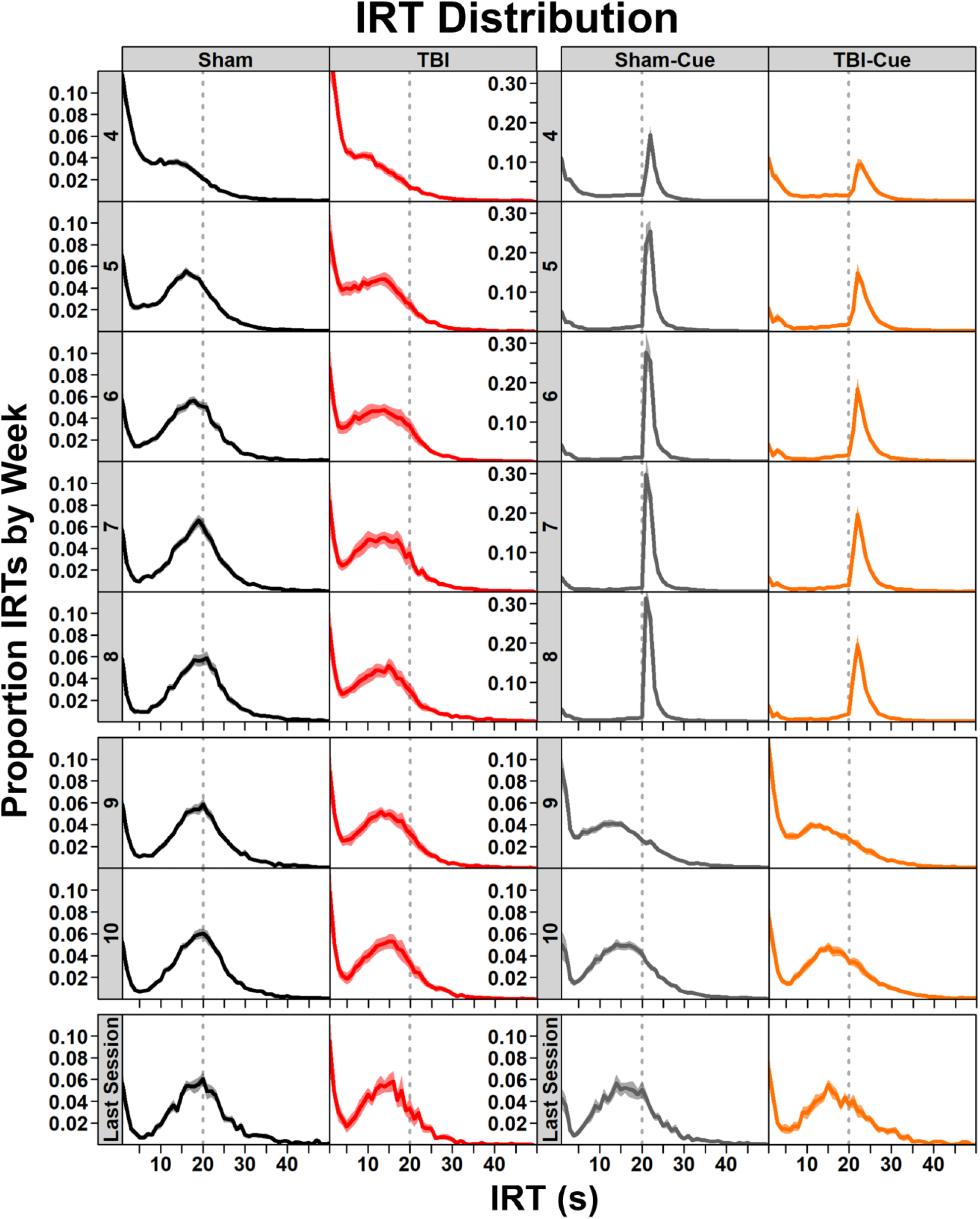
Experiment 3: IRT distributions from the DRL task were used to identify disinhibitory versus timing aspects of impulsivity. Panels from top to bottom show performance across each post-injury week and the final session. The first horizontal break (between week 8 and 9) represents the cessation of cognitive rehabilitation and the second break (after week 10) isolates the last session of post-injury week 10 that was analyzed for persistence of rehabilitative effects. The dotted line indicates the 20 s mark after which presses were reinforced. Note differing Y axes to better display data. Lines represent group means (±SEM shaded).

### Cognitive Rehabilitation Effects Persisted After Discontinuation for TBI Rats

A planned comparison between TBI and TBI-Cue groups was conducted with a t-test for percent reinforced responses, disinhibitory responding AUC, and the timing t_0_ parameter on the final session. There was a significant improvement in the TBI-Cue group in percent reinforced responses (*t*_(17.98)_ = -2.97, *p* = 0.008; Fig 6D) and fewer total presses (*t*_(17.54)_ = -2.46, *p* = 0.025). However, when the variables from the IRT distribution were analyzed, there was no significant improvement in disinhibitory responding (*t*_(18.61)_ = -0.94, *p* = 0.358; Fig 6E), but there were significant improvements in the t_0_ parameter, the ability to accurately time an interval (*t*_(17.58)_ = 2.15, *p* = 0.046; Fig 6F).

## Discussion

There is a need for effective cognitive rehabilitation paradigms to treat individuals suffering from chronic psychiatric symptoms. Animal models for cognitive rehabilitation are necessary to investigate physiological and behavioral mechanisms and optimize therapeutic strategies. Animal models of TBI represent an ideal vehicle for this because the pathophysiological and behavioral features reliably mimic the human condition (Kobeissy et al., 2016). Chronic impulsivity and decision-making deficits occur in patients with brain injuries (Dixon et al., 2005; Moreno-López et al., 2016) and in animal models of injury (Vonder Haar et al., 2016; Shaver et al., 2019). Cognitive rehabilitation paradigms in patients with TBI show some potential (Novakovic-Agopian et al., 2011; Novakovic-Agopian et al., 2021), however, there are also criticisms of the rigor and degree of therapeutic effects for current methods (Fetta et al., 2017). Ultimately, a stronger understanding of the neuroplastic processes underlying these effects could augment therapies through pharmacology, neuromodulation, or modified behavioral paradigms.

In the current study, both behavioral and neurophysiological results suggested outcome insensitivity was underlying behavioral flexibility deficits after frontal TBI. Blunted valence signaling after TBI in frontostriatal brain areas was particularly evident in loci associated with outcome processing such as the lOFC and anterior insula (Ains; Fig 3C). TBI rats demonstrated difficulty adjusting behavior to changing contingencies on the PbR task in line with reward-feedback impairments indicative of poor outcome salience seen at beta frequencies (Fig 2; Fig 3). Therefore, a simple cognitive rehabilitation paradigm to increase the salience of optimal outcomes was evaluated: “correct” responses were cued. This successfully rescued deficits in behavioral flexibility (Fig 4A) and even increased activity of cells in the OFC (Fig 5B), indicating potential for plastic reorganization of these circuits after TBI. We then determined whether this rehabilitation paradigm could translate to a distinct domain of frontostriatal function: impulsivity. Rats with TBI were more impulsive, showing reduced inhibition and waiting during the DRL task (Fig 6; Fig 7). Cognitive rehabilitation also improved these deficits (Fig 6; Fig 7) and, critically, improvement persisted when rehabilitation was discontinued (Fig 6D-F). Together, these data establish a framework for evaluating cognitive rehabilitation efficacy and investigating physiological substrates such as improved outcome salience in reward-related networks.

In experiment 1, local field potentials captured brain dynamics on a mesoscopic scale which offers translational potential to human EEG (Ray et al., 2008; Cui et al., 2016; MacDowell and Buschman, 2020). We identified a reward-feedback signal occurring at beta frequencies that was impaired following TBI. In Sham animals, beta oscillations modulated activity based on reward outcome: higher for rewarded, lower for non-rewarded (Fig 3A,B). This differential was reduced after injury, most prominently in lOFC and Ains, but also in distal regions such as the nucleus accumbens (Fig 3). Beta frequency activity is related to aspects of reward value and outcome evaluation (Pesaran et al., 2008; Torrecillos et al., 2015; Samson et al., 2017; Koloski et al., 2023). In humans, Ains beta oscillations increase during salient outcomes (positive or negative) and correlate with other cortical activity during reward outcome (Haufler et al., 2022). Beta oscillations may be inversely related to reward-prediction error signal (HajiHosseini and Holroyd, 2015), and dopamine disruptions following TBI may modulate changes in beta-driven reward feedback signals. We also observed decreased reward-related delta power following injury (Fig 3A), consistent with a previous report of task-related delta power linked to positive rewards (Cavanagh, 2015). The electrophysiology closely tracked impaired flexibility on the PbR task (Fig 2A-D). TBI rats completed fewer reversals and demonstrated poor initial discrimination suggesting impairments may be driven by outcome evaluation in general. These findings map onto clinical populations with brain injury. The ability to adapt to changing outcomes is impaired after TBI (Schlund, 2002) and patients displayed difficulty identifying the contingencies governing behavioral tasks (Schlund and Pace, 2000). Because of deficits in outcome evaluation, we hypothesized cognitive rehabilitation targeting correct action engagement would be effective. This simple approach is rooted in motor rehabilitation; in constraint-induced movement therapy after stroke, affected limb use is forced by restraining the functional limb (Morris et al., 1997).

We used a simple manipulation for cognitive rehabilitation: cueing correct responses. This substantially improved flexibility (Fig 4A) and impulsivity (Fig 6A). In the PbR task, associating cues with the higher-probability option enabled rats to stay with choices through wins and losses (Fig 4C-D), indicating that TBI rats can use salient signals which may improve physiological mediators of action-outcome associations (Fig 5B). Prior to rehabilitation, there was no injury effect on our 3-session version of the AST (Supplemental Fig 4–1), highlighting the importance of task selection for deficit identification. However, an AST variant using within-session changes (more similar to PbR task) did detect frontal TBI deficits (Craine et al., 2023). In intact rats, reversal learning requires integration of the OFC, PFC, and ventral striatum (Dalton et al., 2014; Dalton et al., 2016; Amodeo et al., 2017). Engagement in cognitive rehabilitation on the PbR task recovered a TBI deficit in OFC activity, as measured by c-Fos staining (Fig 5B), providing further evidence for the OFC in mediating outcome salience deficits. The frontostriatal regions involved in behavioral flexibility are sensitive to changes in dopamine, with impairments from both increased and decreased dopamine activity (Costa et al., 2015; Vo et al., 2016). Interestingly, despite reports on decreased dopamine signaling after TBI (Chen et al., 2015), there was a significant injury-related increase in c-Fos^+^/TH^+^ co-labeled cells in the injury-adjacent PL cortex (Fig 5A).

In the DRL task, using a cue light to indicate the 20 s timer elapsed substantially improved the percent of responses reinforced (Fig 6A), confirming the ability of TBI rats to use salient cues. To better understand this high-resolution behavior, we fit a model to the distribution of IRTs (Fig 1C) and extracted parameters relevant to acute disinhibition (i.e., the area under the initial curve) and ability to accurately time the interval (i.e., t_0_ parameter). Cognitive rehabilitation reduced disinhibitory responding and improved estimation of the time interval (Fig 6B-C, Fig 7). Critically, improvements in TBI rats persisted even after cues were decoupled from reinforcing contingencies (Fig 6D), which was partially mediated by improvements in timing (Fig 6F).

While this is a simplified approach to cognitive rehabilitation, it provides a base to refine clinical applications. The development of effective paradigms is a priority for treating TBI and other neurological and psychiatric disorders. Recent successful studies in patients used similar strategies to the current research: augmenting the salience of outcomes (Dobryakova et al., 2020) or prompting engagement in target behaviors (Rabinowitz et al., 2022). Unfortunately, current solutions are inadequate to the population at large (Kable et al., 2017), highlighting the need for rigorous animal experiments that can also explore neuroplastic mechanisms. Recent animal studies indicate this strategy may be applicable across a variety of injury types and locations. Hippocampal deficits were rescued after a diffuse central brain injury with 7-14 days of free exploration in a complex, changing spatial environment (Law et al., 2023). In a model of lateral, moderate TBI, pre-training on the Morris water maze with no external cues improved performance after injury (Wagner et al., 2013) and post-injury training also improved function (Edwards et al., 2015). Combined with the current study, these data suggest that rehabilitation efforts should be designed to engage specific circuits of interest. A corollary strategy, environmental enrichment, is widely studied in TBI, stroke, and related disorders. Introducing novel toys in larger and/or group housing environment improves motor- and learning-related outcomes after TBI or stroke, though cognitive effects are often less robust than motor (Farrell et al., 2001; Radabaugh et al., 2016; Lajud et al., 2019; Woitke et al., 2023).

Many challenges remain to optimize cognitive rehabilitation for TBI and similar conditions. In the field of stroke, the initiation, duration, and intensity of exercise for rehabilitation is still heavily debated, with cautionary tales against too early *and* too late of intervention (Schmidt et al., 2014; Li et al., 2017). In the current study, we delayed treatment until the chronic period when there are pervasive deficits in impulsivity and decision-making, but no gross deficits in motivation (Vonder Haar et al., 2016; Shaver et al., 2019). The current findings suggest the window for cognitive rehabilitation may be wider than motor (Fig 6D), however this timeframe needs to be defined, and earlier intervention may be more efficacious. Other timing-related approaches such as training on fixed interval schedules reduced impulsive choice in intact rats (Peterson and Kirkpatrick, 2016; Bailey et al., 2018), suggesting there may be many potential strategies to treat impulsive deficits.

The current data provides a proof-of-concept for cognitive rehabilitation in animal models of disease, identifies potential physiological mediators (Fig 2, 3, 5B), and most critically, demonstrates lasting functional recovery (Fig 6D,F). More research will be needed to isolate the physiological substrates and determine how cognitive rehabilitation may be combined with other environmental, pharmacological, or neuromodulatory interventions. While challenging to combine these disparate approaches, basic research will need to incorporate these various strategies to better promote circuit reorganization in models of disease and develop effective treatments for clinical conditions.

## Acknowledgements

We would like to thank Anastasios Lake, Virginia Milleson, Kristen Pechacek, and the other members of the Injury and Recovery Laboratory for helping with behavioral testing as well as Tianzhi Tang, Sidharth Hulyalkar, and Alyssa Terry for help with behavioral training, electrophysiology, and data analysis. This research was funded by the National Institutes for Health (R01-NS110905, P20-GM109098, P30-CA23100, T32-MH018399), VA Office of Research and Development (Career Development Award to DR, Career Development Award IK2BX006125 to MK), VA Excellence for Stress and Mental Health, the Southern Regional Education Board, and West Virginia University.

## Data Availability Statement

Data from experiment 1 will be made publicly available at DANDI: Distributed Archives for Neurophysiology Data Integration (search Dhakshin Ramanathan). Data from experiments 2 and 3 will be made publicly available at the Open Data Commons for Traumatic Brain Injury database (search Vonder Haar). Associated analysis code will be publicly posted to the corresponding author’s Github (https://github.com/VonderHaarLab/).

## Author Contributions

Miranda Koloski: Designed research, performed research, analyzed data, wrote manuscript. Christopher O’Hearn: Designed research, performed research, analyzed data, wrote manuscript. Michelle Frankot: Performed research, analyzed data. Lauren P. Giesler: Performed research, analyzed data. Dhakshin Ramanathan: Designed research, wrote manuscript. Cole Vonder Haar: Designed research, analyzed data, wrote manuscript.

