## Supplemental for "Cognitive rehabilitation can improve brain injury-induced deficits in behavioral flexibility and impulsivity linked to impaired reward-feedback activity"

| | | df | *F* | *p* | | b | *t* | *p* |
|---|---|---|---|---|---|---|---|---|
| **Reversals** | Injury | 1, 22.17 | 3.03 | 0.096 | | | | |
| | Time | 22, 214.48 | 5.31 | **<0.001** | | | | |
| | Injury*Time | 21, 214.49 | 1.82 | **0.001** | Sham v. TBI | -0.22 | -4.24 | **<0.001** |
| **Trials** | Injury | 1, 20.88 | 4.12 | 0.055 | | | | |
| | Time | 22, 213.45 | 3.01 | **<0.001** | | | | |
| | Injury*Time | 21, 213.46 | 1.48 | 0.088 | | | | |
| **Initial Discrimination** | Injury | 1, 4.833 | 8.85 | **0.032** | Sham v. TBI | 0.35 | 4.43 | **<0.001** |
| | Time | 22, 85.78 | 1.05 | 0.421 | | | | |
| | Injury*Time | 21, 85.78 | 0.83 | 0.674 | | | | |
| **p(Switch\|Win)** | Injury | 1, 21.02 | 3.80 | 0.065 | | | | |
| | Time | 22, 244.65 | 4.55 | **<0.001** | | | | |
| | Injury*Time | 21, 244.63 | 1.78 | 0.082 | | | | |
| **p(Switch\|Loss)** | Injury | 1, 20.66 | 5.24 | **0.033** | Sham v. TBI | 0.14 | 2.430 | **0.016** |
| | Time | 22, 247.43 | 2.95 | **0.002** | | | | |
| | Injury*Time | 21, 247.41 | 1.74 | 0.090 | | | | |

**Supplemental Table 2–1.** Statistical parameters from self-paced PbR (Experiment 1) analyses.

| | | Marginal $R^2$ | Cond. $R^2$ | Random Effect Variance |
|---|---|---|---|---|
| **Experiment 1** | **Reversals** | 0.38 | 0.55 | 0.27 |
| Self-paced PbR | **Trials** | 0.27 | 0.54 | 0.36 |
| | **Initial Discrimination** | 0.16 | 0.31 | 0.18 |
| | **p(Switch\|Win)** | 0.19 | 0.33 | 0.01 |
| | **p(Switch\|Loss)** | 0.16 | 0.21 | 0.02 |

**Supplemental Table 2–2.** Individual subject level variance in behavior from Experiment 1. The marginal $R^2$ represents the variance explained by fixed effects. In contrast, the conditional $R^2$ is the variance explained by both the fixed *and* random effects, allowing an indirect measure of individual subject variability. The random effect variance (from subject-level intercept) is reported in units of standard deviation.

| Fixed Effects | | **df** | ***F*** | ***p*** | | **b** | ***t*** | ***p*** |
|---|---|---|---|---|---|---|---|---|
| **lOFC power** | Injury | 1, 60.83 | 3.24 | 0.077 | | | | |
| | Trial | 1, 594.95 | 23.51 | **<0.001** | Reward v. No Reward | 1.67 | 3.24 | **0.001** |
| | Frequency | 5, 58.50 | 3.70 | **0.006** | | | | |
| | Injury*Trial | 1, 594.76 | 43.34 | **<0.001** | Sham v. TBI | -0.11 | -3.01 | **0.003** |
| | Injury*Freq | 5, 58.50 | 0.51 | 0.770 | | | | |
| | Trial*Freq | 5, 594.72 | 13.90 | **<0.001** | | | | |
| | Injury*Trial*Freq | 5, 594.70 | 1.60 | 0.159 | | | | |
| **Beta power** | Injury | 1, 11.20 | 1.13 | 0.310 | | | | |
| | Trial | 1,1329.92 | 208.43 | **<0.001** | Reward v. No Reward | -0.33 | -13.16 | **<0.001** |
| | Electrode | 11, 1325.04 | 3.50 | **<0.001** | | | | |
| | Injury*Trial | 1, 1329.63 | 33.32 | **<0.001** | Sham v. TBI | 0.01 | 0.36 | 0.723 |
| | Injury*Electrode | 11,1325.04 | 2.20 | **0.013** | | | | |
| | Trial*Electrode | 11,1324.97 | 3.25 | **<0.001** | | | | |
| | Injury*Trial*Elec | 11,1324.97 | 1.60 | 0.094 | | | | |

| Random Effects | | **Marginal $R^2$** | **Cond. $R^2$** | **Random Effect Variance** |
|---|---|---|---|---|
| **lOFC Power** | | 0.20 | 0.31 | |
| | **Frequency \| Subject** | | | 0.10 |
| | **Time** | | | 0.03 |
| **Beta Power** | | 0.17 | .047 | |
| | **Subject** | | | 0.33 |
| | **Time** | | | 0.05 |

**Supplemental Table 3–1.** Statistical parameters from multi-channel electrophysiology (Experiment 1) analyses.

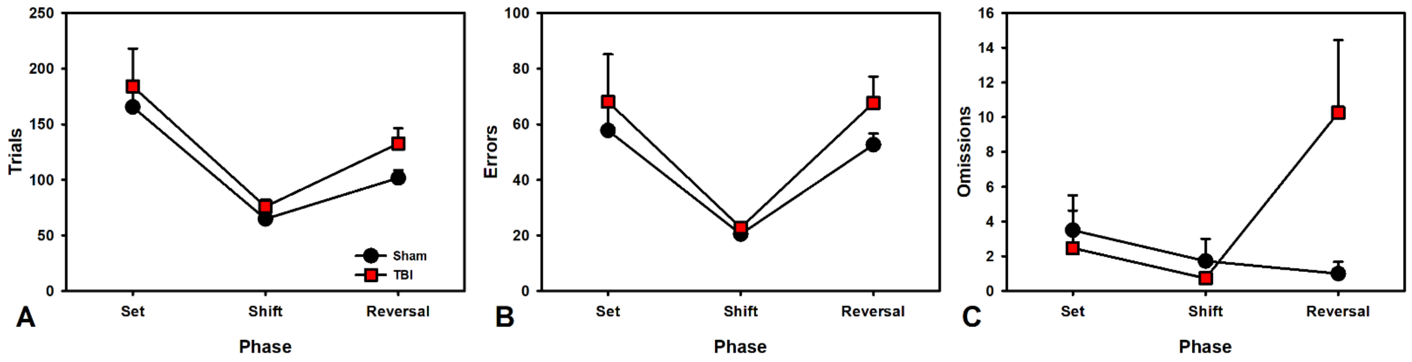


**Supplemental Figure 4–1.** Experiment 2: performance on the AST. A) Trials to criterion across the three phases were analyzed in a 2 X 3 repeated measures ANOVAs [Injury (Sham, TBI) X Phase (Set, Shift, Reversal)]. There was no Injury X Phase interaction ($p$ = 0.851). B) Secondary measures of errors to criterion and omissions were analyzed similarly and revealed no interaction for errors ($p$ = 0.913). C) However, there was a significant interaction for omissions ($p$ = 0.011). Post-hoc analyses using t-tests revealed that TBI rats had significantly more omissions than Sham rats in Phase 3 ($t_{(18.96)}$ = -2.18, $p$ = 0.042).

| | | **df** | ***F*** | ***p*** |
|---|---|---|---|---|
| **Trials** | Injury | 1, 117 | 1.32 | 0.253 |
| | Phase | 2, 117 | 14.32 | **<0.001** |
| | Injury*Phase | 2, 117 | 0.16 | 0.851 |
| **Errors** | Injury | 1, 117 | 0.20 | 0.653 |
| | Phase | 2, 117 | 17.70 | **<0.001** |
| | Injury*Phase | 2, 117 | 0.09 | 0.913 |
| **Omissions** | Injury | 1, 117 | 7.09 | **0.009** |
| | Phase | 2, 117 | 3.81 | **0.025** |
| | Injury*Phase | 2, 117 | 4.72 | **0.011** |
| **Corrects** | Injury | 1, 117 | 1.47 | 0.227 |
| | Phase | 2, 117 | 18.34 | **<0.001** |
| | Injury*Phase | 2, 117 | 0.10 | 0.905 |

**Supplemental Table 4–2.** Statistical parameters from the AST (Experiment 2a) analyses.

| | | df | *F* | *p* | | b | *t* | *p* |
|---|---|---|---|---|---|---|---|---|
| **Switches** | Injury | 1, 37.16 | 0.00 | 0.999 | | | | |
| | Rehab | 1, 37.16 | 82.71 | **<0.001** | Non-Cued v. Cued | 1.02 | 5.65 | **<0.001** |
| | Week | 1, 921.28 | 325.76 | **<0.001** | | | | |
| | Injury*Rehab | 1, 37.16 | 1.92 | 0.174 | | | | |
| | Injury*Week | 1, 921.28 | 0.96 | 0.327 | | | | |
| | Rehab*Week | 1, 921.28 | 117.76 | **<0.001** | Non-Cued v. Cued | 0.49 | 9.45 | **<0.001** |
| | Injury*Rehab*Week | 1, 921.28 | 4.16 | **0.042** | Sham v. TBI | 0.11 | 2.23 | **0.026** |
| | | | | | Sham-Cue v. TBI-Cue | -0.04 | -0.72 | 0.473 |
| **Omissions** | Injury | 1, 36.98 | 1.69 | 0.201 | | | | |
| | Rehab | 1, 36.98 | 0.14 | 0.713 | | | | |
| | Week | 1, 921.04 | 17.56 | **<0.001** | | | | |
| | Injury*Rehab | 1, 36.98 | 0.46 | 0.502 | | | | |
| | Injury*Week | 1, 921.04 | 33.74 | **<0.001** | Sham v. TBI | -0.18 | -2.99 | **0.003** |
| | Rehab*Week | 1, 921.04 | 0.01 | 0.926 | | | | |
| | Injury*Rehab*Week | 1, 921.04 | 3.13 | 0.077 | | | | |
| **p(Switch\|Win)** | Injury | 1, 37.15 | 0.89 | 0.350 | | | | |
| | Rehab | 1, 37.15 | 36.46 | **<0.001** | Non-Cued v. Cued | 0.74 | 3.39 | **0.002** |
| | Week | 1, 921.23 | 424.85 | **<0.001** | | | | |
| | Injury*Rehab | 1, 37.15 | 1.99 | 0.167 | | | | |
| | Injury*Week | 1, 921.23 | 4.21 | **0.041** | Sham v. TBI | -0.15 | -2.65 | **0.008** |
| | Rehab*Week | 1, 921.23 | 87.55 | **<0.001** | Non-Cued v. Cued | 0.43 | 8.21 | **<0.001** |
| | Injury*Rehab*Week | 1, 921.23 | 3.42 | 0.065 | | | | |
| **p(Switch\|Loss)** | Injury | 1, 36.9 | 6.95 | **0.012** | Sham v. TBI | 0.45 | 2.27 | **0.029** |
| | Rehab | 1, 36.9 | 8.52 | **0.006** | Non-Cued v. Cued | 0.49 | 2.48 | **0.018** |
| | Week | 1, 921.09 | 63.97 | **<0.001** | | | | |
| | Injury*Rehab | 1, 36.9 | 0.21 | 0.650 | | | | |
| | Injury*Week | 1, 921.09 | 0.84 | 0.360 | | | | |
| | Rehab*Week | 1, 921.09 | 97.30 | **<0.001** | Non-Cued v. Cued | 0.60 | 8.44 | **<0.001** |
| | Injury*Rehab*Week | 1, 921.09 | 2.72 | 0.099 | | | | |
| **Percent Correct** | Injury | 1, 36.98 | 1.09 | 0.303 | | | | |
| | Rehab | 1, 36.98 | 177.12 | **<0.001** | Non-Cued v. Cued | 1.04 | 7.56 | **<0.001** |
| | Week | 1, 921.15 | 413.66 | **<0.001** | | | | |
| | Injury*Rehab | 1, 36.98 | 8.96 | **0.005** | Sham v. TBI | -0.20 | -1.43 | 0.161 |
| | | | | | Sham-Cue v. TBI-Cue | 0.41 | 2.76 | **0.009** |
| | Injury*Week | 1, 921.15 | 2.20 | 0.138 | | | | |
| | Rehab*Week | 1, 921.15 | 220.01 | **<0.001** | Non-Cued v. Cued | 0.58 | 12.16 | **<0.001** |
| | Injury*Rehab*Week | 1, 921.15 | 3.07 | 0.080 | | | | |
| **Reinforcers** | Injury | 1, 37.1 | 0.16 | 0.689 | | | | |
| | Rehab | 1, 37.1 | 78.47 | **<0.001** | Non-Cued v. Cued | 0.89 | 4.95 | **<0.001** |
| | Week | 1, 921.22 | 269.56 | **<0.001** | | | | |
| | Injury*Rehab | 1, 37.1 | 4.44 | **0.042** | Sham v. TBI | -0.22 | -1.25 | 0.219 |
| | | | | | Sham-Cue v. TBI-Cue | 0.33 | 1.71 | 0.095 |
| | Injury*Week | 1, 921.22 | 0.57 | 0.451 | | | | |
| | Rehab*Week | 1, 921.22 | 104.70 | **<0.001** | Non-Cued v. Cued | 0.46 | 8.54 | **<0.001** |
| | Injury*Rehab*Week | 1, 921.22 | 2.02 | 0.155 | | | | |
| **Collection Latency** | Injury | 1, 37.17 | 22.29 | **<0.001** | Sham v. TBI | 1.20 | 4.06 | **<0.001** |
| | Rehab | 1, 37.17 | 0.00 | 0.976 | | | | |
| | Week | 1, 848.93 | 13.53 | **<0.001** | | | | |
| | Injury*Rehab | 1, 37.17 | 0.66 | 0.423 | | | | |
| | Injury*Week | 1, 848.93 | 0.96 | 0.326 | | | | |
| | Rehab*Week | 1, 848.93 | 0.36 | 0.549 | | | | |
| | Injury*Rehab*Week | 1, 848.93 | 0.02 | 0.894 | | | | |

**Supplemental Table 4–3.** Statistical parameters from PbR (Experiment 2b) analyses.

| | | Marginal $R^2$ | Cond. $R^2$ | Random Effect Variance |
|---|---|---|---|---|
| **Experiment 2** PbR | **Switches** | 0.50 | 0.66 | 0.40 |
| | **Omissions** | 0.13 | 0.94 | 1.69 |
| | **p(Switch\|Win)** | 0.42 | 0.66 | 0.50 |
| | **p(Switch\|Loss)** | 0.19 | 0.37 | 0.43 |
| | **Percent Correct** | 0.63 | 0.72 | 0.30 |
| | **Reinforcers** | 0.48 | 0.64 | 0.40 |
| | **Collection Latency** | 0.28 | 0.74 | 0.68 |

**Supplemental Table 4–4.** Individual subject level variance in behavior from Experiment 2. The marginal $R^2$ represents the variance explained by fixed effects. In contrast, the conditional $R^2$ is the variance explained by both the fixed *and* random effects, allowing an indirect measure of individual subject variability. The random effect variance (from subject-level intercept) is reported in units of standard deviation, with the exception of Omissions which are raw values from count data under a Poisson GLMER.

| | | c-Fos+ | | | c-Fos+/TH+ | | |
|---|---|---|---|---|---|---|---|
| | | **b** | ***z*** | ***p*** | **df** | ***F*** | ***p*** |
| **PL** | Injury | 0.72 | 1.39 | 0.165 | 1,21 | 4.85 | **0.039** |
| | Rehab | 0.42 | 0.75 | 0.456 | 1,21 | 0.00 | 0.989 |
| | Injury*Rehab | -1.28 | -1.50 | 0.134 | 1,21 | 0.81 | 0.379 |
| **OFC** | Injury | 0.46 | 1.35 | 0.176 | 1,24 | 0.03 | 0.870 |
| | Rehab | 0.89 | 2.33 | **0.020** | 1,24 | 1.11 | 0.302 |
| | Injury*Rehab | -1.58 | -2.73 | **0.006** | 1,24 | 0.00 | 0.954 |
| **NAC** | Injury | 0.25 | 0.66 | 0.510 | 1,19 | 0.06 | 0.812 |
| | Rehab | 0.71 | 1.66 | 0.096 | 1,19 | 0.01 | 0.916 |
| | Injury*Rehab | -1.05 | -1.61 | 0.107 | 1,19 | 1.65 | 0.214 |

**Supplemental Table 5–1.** Statistical parameters from immunohistochemistry (Experiment 2c) analyses.

| | | df | *F* | *p* | | b | *t* | *p* |
|---|---|---|---|---|---|---|---|---|
| **Percent Reinforced** | Injury | 1,40.99 | 20.68 | **<0.001** | Sham v. TBI | -0.65 | -5.40 | **<0.001** |
| | Rehab | 1,40.99 | 382.17 | **<0.001** | Non-Cued v. Cued | 1.42 | 12.07 | **<0.001** |
| | Week | 1,1013.34 | 588.26 | **<0.001** | | | | |
| | Injury*Rehab | 1,40.99 | 9.23 | **0.004** | Sham v. TBI | 0.52 | 3.04 | **0.004** |
| | Injury*Week | 1,1013.34 | 38.76 | **<0.001** | Sham v. TBI | 0.35 | 17.72 | **<0.001** |
| | Rehab*Week | 1,1013.34 | 4.15 | **0.042** | Non-Cued v. Cued | -0.05 | -1.87 | 0.061 |
| | Injury*Rehab*Week | 1,1013.34 | 21.19 | **<0.001** | Sham v. TBI | -0.23 | -7.68 | **<0.001** |
| | | | | | Sham-Cue v. TBI-Cue | -0.03 | -1.14 | 0.253 |
| **Presses** | Injury | 1,41 | 4.08 | **0.050** | Sham v. TBI | -0.76 | -3.63 | **0.001** |
| | Rehab | 1,41 | 62.40 | **<0.001** | Non-Cued v. Cued | 0.72 | 3.52 | **0.001** |
| | Week | 1,1013.24 | 566.37 | **<0.001** | | | | |
| | Injury*Rehab | 1,41 | 9.48 | **0.004** | Sham v. TBI | 0.92 | 3.08 | **0.004** |
| | Injury*Week | 1,1013.24 | 5.40 | **0.020** | Sham v. TBI | 0.47 | 16.78 | **<0.001** |
| | Rehab*Week | 1,1013.24 | 1.72 | 0.191 | | | | |
| | Injury*Rehab*Week | 1,1013.24 | 14.83 | **<0.001** | Sham v. TBI | -0.19 | -4.38 | **<0.001** |
| | | | | | Sham-Cue v. TBI-Cue | 0.05 | 1.08 | 0.282 |
| **Disinhibition (AUC)** | Injury | 1,41 | 1.14 | 0.292 | | | | |
| | Rehab | 1,41 | 0.18 | 0.672 | | | | |
| | Week | 1,176 | 114.05 | **<0.001** | | | | |
| | Injury*Rehab | 1,41 | 0.08 | 0.779 | | | | |
| | Injury*Week | 1,176 | 0.04 | 0.846 | | | | |
| | Rehab*Week | 1,176 | 6.61 | **0.011** | Non-Cued v. Cued | -0.24 | -2.29 | **0.023** |
| | Injury*Rehab*Week | 1,176 | 0.31 | 0.577 | | | | |
| **Timing (t0)** | Injury | 1,41 | 6.97 | **0.005** | Sham v. TBI | -0.74 | -6.36 | **<0.001** |
| | Rehab | 1,41 | 409.10 | **<0.001** | Non-Cued v. Cued | 1.18 | 10.42 | **<0.001** |
| | Week | 1,176 | 13.32 | **<0.001** | | | | |
| | Injury*Rehab | 1,41 | 33.09 | **<0.001** | Sham v. TBI | 0.99 | 5.99 | **<0.001** |
| | Injury*Week | 1,176 | 9.53 | **0.007** | Sham v. TBI | 0.47 | 9.76 | **<0.001** |
| | Rehab*Week | 1,176 | 31.01 | **<0.001** | Non-Cued v. Cued | -0.53 | -7.43 | **<0.001** |
| | Injury*Rehab*Week | 1,176 | 9.93 | **0.006** | Sham v. TBI | -0.29 | -3.92 | **<0.001** |
| | | | | | Sham-Cue v. TBI-Cue | 0.00 | 0.04 | 0.968 |

**Supplemental Table 6–1.** Statistical parameters from DRL (Experiment 3a) analyses.

| | | **Marginal $R^2$** | **Cond. $R^2$** | **Random Effect Variance** |
|---|---|---|---|---|
| **Experiment 3** DRL | **Percent Reinforced** | 0.80 | 0.88 | 0.28 |
| | **Presses** | 0.53 | 0.76 | 0.49 |
| | **Disinhibition (AUC)** | 0.19 | 0.67 | 0.71 |
| | **Timing (t0)** | 0.77 | 0.80 | 0.18 |

**Supplemental Table 6–2.** Individual subject level variance in behavior from Experiment 3. The marginal $R^2$ represents the variance explained by fixed effects. In contrast, the conditional $R^2$ is the variance explained by both the fixed *and* random effects, allowing an indirect measure of individual subject variability. The random effect variance (from subject-level intercept) is reported in units of standard deviation.
